# Kinetics of *Plasmodium midgut* invasion in Anopheles mosquitoes

**DOI:** 10.1101/2020.02.06.937276

**Authors:** Gloria Volohonsky, Perrine Paul-Gilloteaux, Jitka Štáfková, Julien Soichot, Jean Salamero, Elena A. Levashina

## Abstract

Malaria-causing *Plasmodium* parasites traverse the mosquito midgut cells to establish infection at the basal side of the midgut. This dynamic process is a determinant of mosquito vector competence, yet the kinetics of the parasite migration is not well understood. Here we used transgenic mosquitoes of two *Anopheles* species and a *Plasmodium berghei* fluorescence reporter line to track parasite passage through the mosquito tissues at high spatial resolution. We provide new quantitative insight into malaria parasite invasion in African and Indian *Anopheles* species and demonstrate that species-specific kinetics of *Plasmodium* invasion is shaped by the mosquito complement-like system.

**Author Summary:** The traversal of the mosquito midgut cells is one of the critical stages in the life cycle of malaria parasites. Motile parasite forms, called ookinetes, traverse the midgut epithelium in a dynamic process which is not fully understood.

Here, we harnessed transgenic reporters to track invasion of *Plasmodium* parasites in African and Indian mosquito species. We found important differences in parasite dynamics between the two anopheline species and demonstrated an unexpected role of mosquito complement-like system in regulation of parasite invasion.

## Introduction

Malaria is a vector-borne human infectious disease caused by protozoan parasites of *Plasmodium* species. It is widespread in tropical and subtropical regions, including parts of the Americas, Asia, and Africa. Approximately 200 million annual cases of malaria result in half a million deaths [1]. Malaria-causing *Plasmodium* parasites are transmitted by *Anopheline* mosquitoes. Among more than 400 of known *Anopheles* species, only 40 are vectors of human malaria [2].

*Plasmodium* development in the mosquito begins with the ingestion of red blood cells infected with sexual-stage gametocytes. In the mosquito midgut, gametocytes differentiate into gametes that egress from the red blood cells and fuse to form the zygotes that develop into motile ookinetes within 16-18 h. The ookinetes penetrate the midgut epithelium 18 – 26 h after the infectious blood meal and transform into vegetative oocysts on the basal side of the midgut [3]. After 12–14 days, mature oocysts rupture and release thousands of sporozoites into the mosquito hemocoel. Released sporozoites invade the salivary glands, where they reside inside the salivary ducts to be injected into a new host when the infected mosquito feeds again [4].

The passage of the malaria parasite through the mosquito vector is characterized by a major population bottleneck. Previous studies revealed that mosquitoes kill the majority of invading *Plasmodium* parasites (reviewed by [5,6]), predominantly during the ookinete stage at the basal side of the epithelium [7].

The immune response of mosquitoes to *Plasmodium* parasites is multifaceted and involves multiple processes. In the midgut, reactive oxygen and nitrogen species, hemoglobin degradation products, as well as digestive enzymes and ‘bacterial flora, all affect the rate of *Plasmodium* development (Reviewed in [8]). As parasites traverse midgut epithelial cells, the invaded cells produce high levels of nitric oxide synthase and peroxidases, creating a toxic environment for the parasites. As a result, some parasites undergo nitration which marks them for killing by the mosquito complement-like system [9]. Furthermore, intracellular parasites can trigger apoptosis causing extrusion and clearance of invaded cells from the cellular layer into the midgut lumen [10]. As *Plasmodium* tries to evade reactive oxygen and nitrogen species inside the cells, these toxic molecules may shape the path taken by the parasite through the cellular layer. When the surviving parasites finally reach the basal lamina, they encounter soluble immune factors that circulate in the hemolymph. Complement-like proteins TEP1 and leucine-rich repeat proteins APL1 and LRIM1 form a complex that mediates parasite killing [11,12]. Histological studies have shown that parasites crossing the cellular layer can be found both inside and in between midgut cells [3,13]. However, it is not yet known whether some parasites cross the cellular layer exclusively between cells, thus avoiding nitration and subsequent recognition by TEP1.

Despite accumulating evidence of molecular processes that govern the passage of motile ookinetes through mosquito tissues, the complexity and diversity of this dynamic process remains to be deciphered. Three modes of motility were reported for the invading ookinetes, namely spiraling, gliding and stationary rotation [14][15]. Spiraling and gliding movements result in active displacement of the parasite in space. In contrast, stationary rotation movement was observed for prolonged periods of time and resulted in no displacement of the ookinete. Because of the lack of markers of the entire midgut cellular layer, previous studies did not establish how distinct types of movements correlate with ookinete location in the midgut.

It has been previously demonstrated that *Anopheles* species differ in their vector competence [16]. In the laboratory, *Anopheles stephensi* (*As*) and *Anopheles gambiae (Ag)* can be infected with the murine parasite *Plasmodium berghei*, albeit at different rates [17]. We set out to image *in vivo* migration of the RFP-expressing *P. berghei* (Pb) ookinetes through the epithelial cells in these two genetically-modified mosquito species that express GFP in the midgut cells. Using high-speed spinning disk microscopy and automated image analyses, we quantified parasite invasion dynamics at high spatial and temporal resolution. Our data revealed unexpected differences in invasion of closely-related mosquito species, pointing to important species-specific mechanisms that regulate mosquito – parasite interactions. Moreover, silencing of the major component of the mosquito complement-like system affected the parasite invasion dynamics, suggesting a new function of TEP1 at the early stages of the midgut invasion process.

## Results and discussion

### TEP1 inhibits midgut invasion of *P. berghei* ookinetes

To study the passage of *Pb* ookinetes through the mosquito midgut, we combined multiscale imaging techniques with high-throughput data analysis and mining (Fig 1). We used transgenic mosquitoes expressing GFP under midgut-specific promoters [18,19] to label mosquito midgut cells, and transgenic rodent *Pb* parasites expressing RFP under a constitutive promoter [20] (S1a Fig). We first made sure that expression of the reporters did not interfere with *Plasmodium* infection. As expected, a significant difference was observed in infection intensity between *As* and *Ag*. Regardless of the infection levels, *As* developed significantly higher oocysts numbers than *Ag* (S1b Fig). We concluded that the transgenic mosquito and *Pb* lines can be used for *in vivo* imaging.

**Figure 1.**
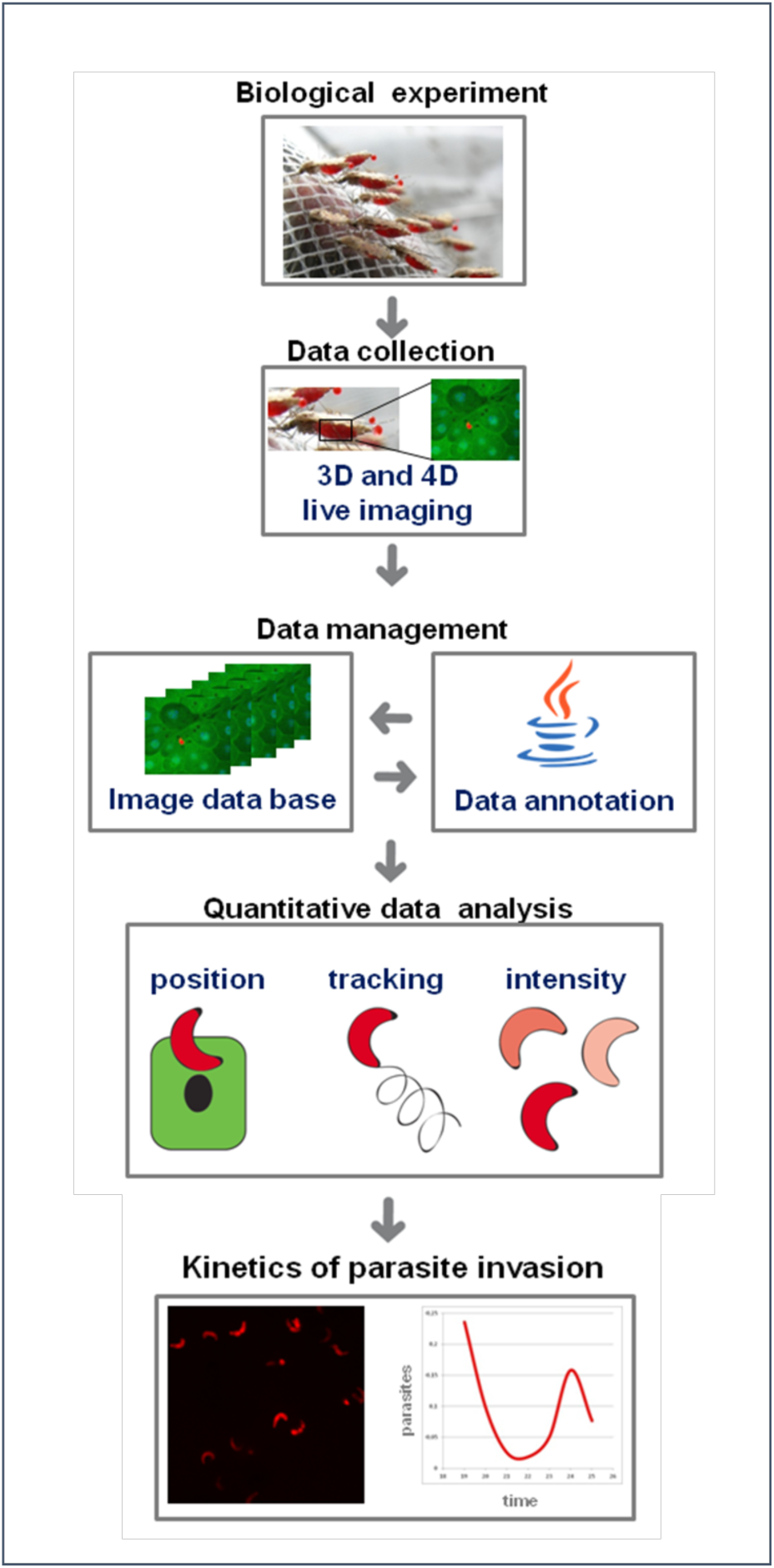
Workflow and experimental settings. *A. stephensi* (*As*) and *A. gambiae* (*Ag*) mosquitoes were blood fed on *P. berghei* infected mice, their midguts dissected and visualized using fast confocal microscopy. Images from all experiments collected at different time points after infection were uploaded into an image database and annotated. Quantitative data was extracted from the images in the database regarding the number, position and intensity of visualized parasites. The results of the data analysis reveal the kinetics of parasite invasion.

As the transgenic mosquito lines expressed GFP in the entire midgut cell, we measured the exact position of RFP-expressing parasites relative to the cellular layer (Fig 2). To this end, we collected large series of z-stack images of live parasites inside the dissected mosquito midguts at different time points after infection and time-lapse images of selected parasites. These tools enabled us to study the parasite invasion process at two time-scales: one was based on statistical analysis of parasites in three dimensional (3D) snapshots of the state of infection between 18 and 25 h post infection, the second tracked single parasites 18 to 25 h post infection over a time of 20 min to 2 h.

**Figure 2.**
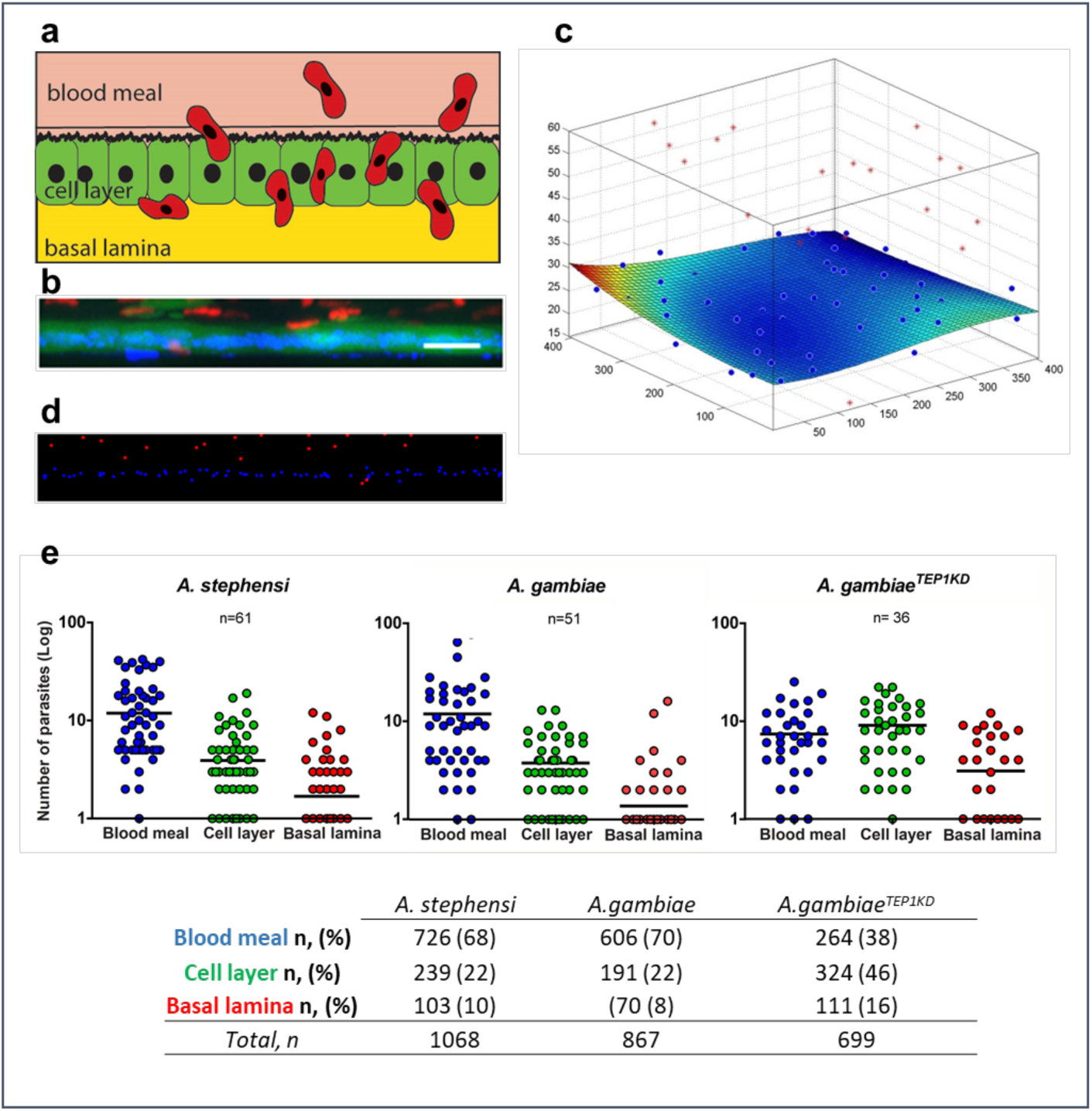
Positions of the parasites relative to the midgut cells. **a**. Schematic representation of topology in the mosquito midgut. Motile ookinetes (red) traverse the mosquito midgut cells and establish infection on the basal side under the basal lamina. **b**. A representative projection of a cross section of *A. stephensi* midgut, scale bar - 50 µm. GFP-positive midgut cells are in green, RFP-positive *P. berghei* parasites are in red, nuclei are labeled by DAPI in blue. **c**. Schematic 3D representation of the same midgut as in (**b)**, where the position of the cellular layer is calculated relative to the nuclei. Positions of parasites are indicated as red dots, nuclei as blue dots. Deviation of the cell layer from a flat surface is color-coded from blue to red (blue no deviation, red - 10 µm). Note the blood meal location of the majority of parasites (above the cell layer). **d**. Representation of nuclei (blue) and parasites (red) in the same midgut as (**b)** after segmentation. **e**. Pooled positions of the parasites from all records at all time points are shown for three layers relative to the midgut cells (blood meal, cellular layer or basal lamina) for *A. stephensi, A. gambiae* and *A. gambiae* mosquitoes silenced for *TEP1* (*A. gambiae*^*TEP1KD*^). Each dot represents the number of parasites at a given position in a single midgut. The numbers of midguts analyzed (n) are indicated above the graph. Horizontal lines depict the mean number of parasites per position. The table below summarizes parasite distribution inside the mosquito midguts at 18-25 hpi. The percentage of ookinetes in the midguts of *A. stephensi, A. gambiae* and *Ag*^*TEP1KD*^ at each location (blood meal, cellular layer and basal lamina) is given in parenthesis. n is the number of parasites at each position, total n is the total number of analyzed parasites.

For each record, parasites and nuclei of the midgut cells were segmented and their positions in 3D space were calculated relative to the cellular layer at each examined time point after infection (Fig 2). The position of parasites relative to the cellular layer was determined by fitting the midgut cell nuclei position by a cubic spline surface. This surface was then considered as the central position of the cellular layer (normalized z=0). An average thickness of 5 µm above and below this surface defined the average cellular layer position.

We next examined whether the dynamics of parasite invasion was similar in two *Anopheles* species. To this end, we measured the number of parasites at each position (blood meal, cellular layer and basal lamina) in *As* and *Ag*. Analyses of all time points did not detect significant differences in parasite localization between the two species (Fig 2e). The majority of ookinetes were detected in the blood meal (70%) and in the cellular level (20%). Only few ookinetes crossed the midgut and reached the basal side (10%). Interestingly, silencing of the major antiparasitic factor *TEP1* in *Ag* (*AgTEP1*^*KD*^) significantly changed spatial distribution of the parasites with only 40% of ookinetes observed in the blood meal, 45% in the cellular layer and 15% at the basal side. This difference in the dynamics of *Pb* invasion in *TEP1*-depleted mosquitoes was suggestive of an additional role of TEP1 in inhibition of ookinete midgut invasion. Previous studies reported *TEP1* expression in the larval gastric ceaca and adult midguts [21,22]. In line with these reports, silencing of *TEP1* also affected midgut microbiota by an as yet unknown mechanism [23]. Our findings extend these observations to the early stages of parasite invasion and suggest that in addition to parasite killing at the basal side, TEP1 directly or indirectly inhibits *Plasmodium* midgut traversal.

### Dynamics of the ookinete midgut invasion

We next focused on *P. berghei* ookinete passage through the mosquito midgut cells at different time points after infection and examined the proportion of parasites at each position (blood meal, cellular layer and basal lamina). To this end, we calculated the average proportion of parasites at each position at the early (18 – 20 h post infection, hpi), intermediate (21 -23 hpi) and late (24 – 25 hpi) intervals after infection (Fig 3a, S4Fig).

**Figure 3.**
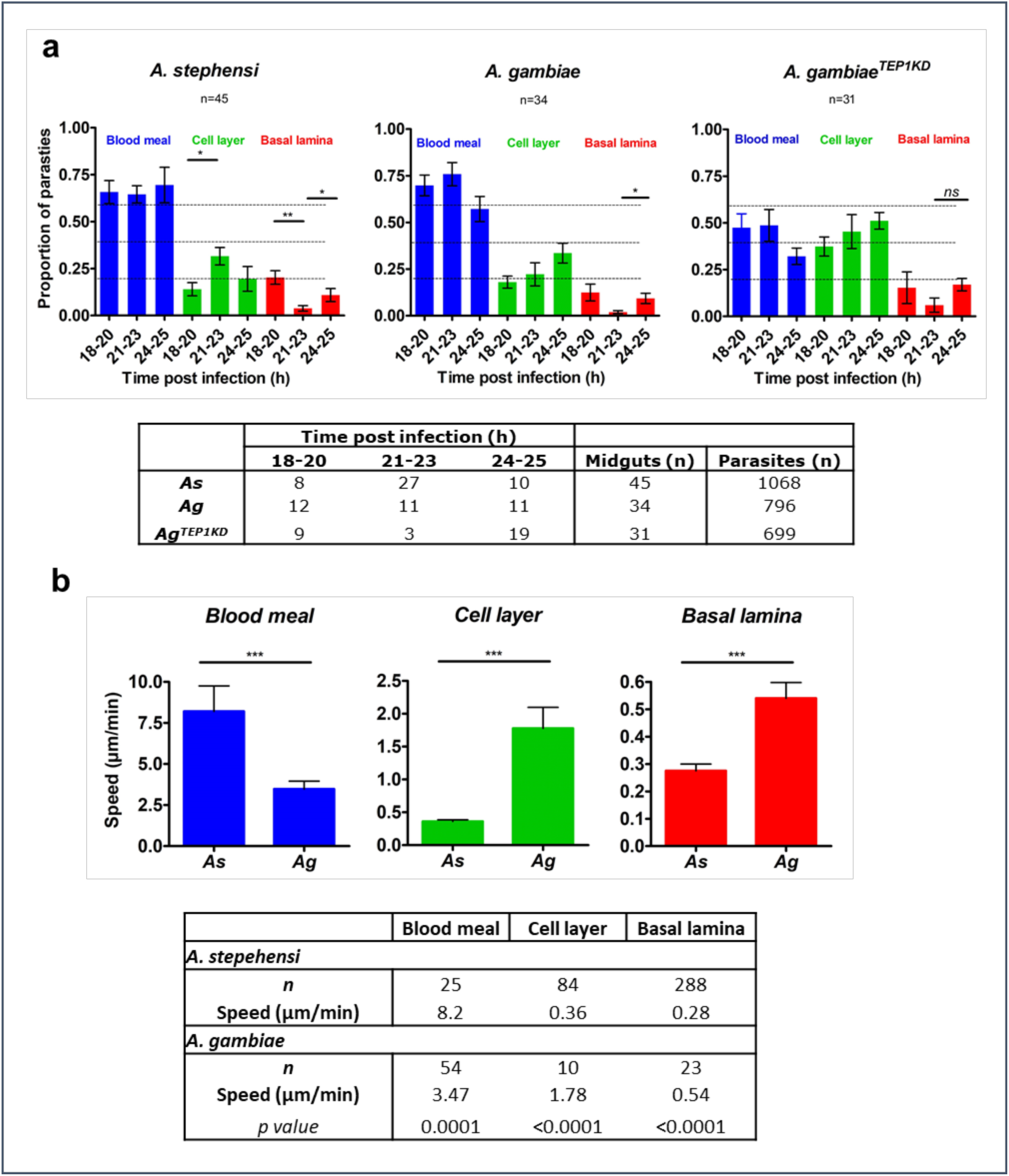
Kinetics of *P. berghei* invasion of *A. stephensi* and *A. gambiae* midguts. **a.** Positions of parasites in *A. stephensi* (*As*), *A. gambiae* (*Ag*) and in *A. gambiae* mosquitoes silenced for *TEP1* (*A. gambiae*^*TEP1KD*^) between 18 and 25 h post infection (hpi). Plots show the proportion of parasites at each position (blood meal, cellular layer and basal lamina) for three different time intervals (18-20, 21-23 and 24-25 hpi). Each bar represents the average proportion of parasites in midguts with at least 10 parasites. Parasite positions were calculated by the distance from the cellular layer: blood meal for ookinetes detected more than 5 µm above the cellular layer; basal lamina for parasites observed more than 5 µm below the cellular layer. Statistical analyses were performed by a non-parametric Mann-Whitney test. The table below shows the number of midguts analyzed at each time interval for each mosquito type. **b.** Speed of parasites as function of the parasite position in *As* and *Ag*. Speed (µm/min) was determined by tracking the parasites position over time from the time-lapse series. Four time-lapse experiments were used: guid 1615 and guid 1628 for *As* and guid 1622 and guid 2109 for *Ag*. The table below details the number of frames (n) used for speed calculations. Statistical significance of differences in the average speed at each given position between *As* and *Ag* were examined by the non-parametric Mann-Whitney test and *p* ≤ 0.0001 are shown by three asterisks.

We observed that in *As* mosquitoes the proportion of blood bolus-residing parasites did not change over time. The proportion of parasites within the cellular layer significantly increased between the early (14% at 18-20 hpi) and intermediate (32% at 21-23 hpi) time intervals. However, this increase did not cause accumulation of the ookinetes at the basal lamina. Instead, a significant decrease in the proportion of basally located parasites was detected between the early (20% at 18-20 hpi) and intermediate (4% at 21-23 hpi) time intervals. Strikingly, this decrease was temporal, as the proportion of parasites in the basal lamina significantly increased at the late time interval (10% at 24-25 hpi). Similar decrease in the proportion of basally located ookinetes was detected in *Ag*, where the proportion of parasites at the basal lamina declined from 12% at 18-20 hpi to 3% at 21-23 hpi, and then increased again to 14% at the late time interval.

Since the mosquito immune system targets the ookinetes at the basal side of the midgut [24], we examined whether the observed decrease in the proportion of basally located ookinetes was rescued by *TEP1* knockdown. *TEP1* silencing eliminated the decrease in the basally located ookinetes observed in *As* and *Ag* mosquitoes and at the same time increased the proportion of parasites within the cellular layer (Fig 3a). These results suggest that the first wave of invading ookinetes is rapidly killed and lysed by the mosquito immune system. As the parasites that reach the basal lamina at later time points do accumulate, it is possible that asynchronous midgut invasion by *Pb* exhausts the components of the mosquito immune system and, thereby, benefits the establishment of infection by the second wave of the parasites. These results may also explain why not all parasites are recognized and killed by TEP1 at the basal lamina. We suggest that early crossing parasites may serve as pioneers that attract and locally deplete TEP1, allowing later-coming parasites to survive the immune attack.

To better understand *Pb* invasion dynamics, we measured ookinete motility in time-lapse experiments. The blood-filled midguts were dissected from infected mosquitoes and mounted *ex vivo* for imaging by spinning disk microscopy for 20 to 120 min. In line with the previous work [14], we observed four distinct ookinete motility modes: (i) passive floating within the blood bolus (guid 2107, guid 1615, S1 Table), (ii) gliding within the cellular layer (guid 1628, S1 Table) (iii) spiraling in the blood meal and within the cellular layer (guid 1622, guid 1624, S2 Table) and (iv) stationary rotation without translocation within the cellular layer (guid 2115, S1 Table). Some ookinetes were observed within a midgut cell for more than one hour, suggesting that the parasites may remain intracellular for relatively long periods of time without inducing cellular apoptosis. By measuring the parasite speed in the blood meal, cellular layer, and at the basal lamina, we found that the speed of ookinetes carried by the bolus content was the highest as compared to other locations (Fig 3b). Interestingly, the speed of the ookinetes in the blood bolus differed between *As* (8.2 µm/min) and *Ag* (3.4 µm/min) midguts, suggesting some differences in the blood bolus environment. The ookinete spiraling motility in the cellular layer was much slower in both mosquito species, namely 0.36 µm/min in *As* and 1.78 µm/min in *Ag*. The slowest stationary rotation movement of parasites was observed at the basal lamina (in *As*, average speed 0.28 µm/min, guid 2113, S1 Table, in *Ag*, average speed 0.54 µm/min, guid 1622, S2 Table). We noted that the speed of ookinetes within the cellular layer and at the basal lamina was faster in *Ag* mosquitoes than in *As* mosquitoes. This observation indicates important differences in the cellular organization of midguts of the closely related mosquito species.

### Ookinete invasion routes

To characterize ookinete invasion routes, intra- or extracellular location of the ookinetes at the cellular layer was examined in more detail. To this end, we developed an algorithm that classified intracellular, extracellular and intercellular parasites based on the score of their 3D distance to the four nearest neighboring nuclei of the midgut cells. The score was calculated for each parasite (Fig 4a,b). The parasites with the score between 0 - 0.45 were defined as extracellular, 0.45-0.55 - as intercellular, and higher than 0.55 - as intracellular. We noticed a proportion of parasites that was extracellular at all time points in both species (Fig 4c).

**Figure 4.**
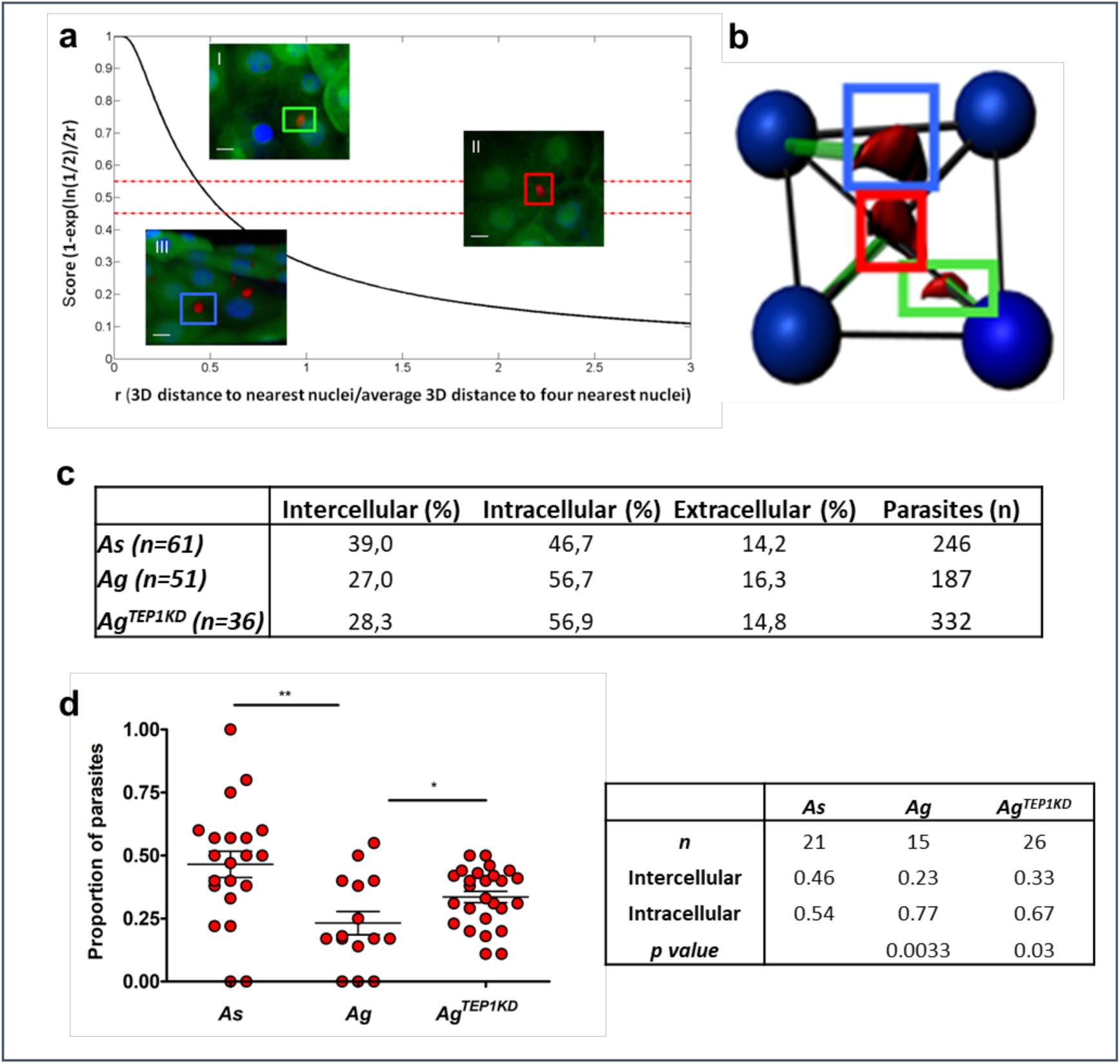
Parasite distribution in the mosquito midgut. Parasite positions within the cellular layer calculated relative to the distance of each parasites to the nuclei of surrounding midgut cells. **a**. Calculations of the distance of parasites from the nuclei of the nearest neighboring midgut cell. The score (s) determines whether the parasite is intercellular (0.45 ≤ s ≤ 0.55), extracellular (s < 0.45), or intracellular (s>0.55). Example images from a z stack, scale bar = 20 µm: (I) s = 0.74, the parasite (green arrow) is intracellular; (II) s = 0.45 (red arrow) the parasite is intercellular and (III) s = 0.36, the parasite is extracellular (blue arrow). **b**. Schematic representation of parasite (red) and nuclei (blue) positions with distances (green lines) used to calculate distances from the nuclei. **c**. Positions of parasites within the cell layer over time in *A. stephensi* (*As*), *A. gambiae* (*Ag*) and *A. gambiae* mosquitoes silenced for *TEP1* (*A. gambiae*^*TEP1KD*^). The table indicates the percentage of parasites at each position for each mosquito. The number *(n)* indicates the number of midguts analyzed for each mosquito genotype. **d.** Comparison of the proportion of intercellular parasites between *As, Ag* and *Ag*^*TEP1KD*^. Each dot represents the proportion of parasites detected between cells in a single midgut. Midguts (n) with at least six parasites within the cellular layer were used for analyses. Statistically significant differences between *As* and *Ag* and between *Ag* and *Ag*^*TEP1KD*^ revealed by a non-parametric t-test (Mann-Whitney) are indicated by asterisks (*p* = 0.03 (*); *p* = 0.003(**)). The table details the mean proportion values for parasites in each midgut and for each position for *n* mosquitoes.

When comparing intercellular and intracellular parasite distribution, a higher proportion of intercellular ookinetes was observed in *As* (40%) than in *Ag* (20%) (Fig 4d, S6 Fig). These results point to intricate differences in parasite invasion routes between the two related *Anopheline* species.

### Parasite viability within the midgut

As the transgenic *P. berghei* line used in this study expressed the fluorescence reporter under a constitutive promoter, we were surprised by high variability in the reporter fluorescence levels observed between individual parasites in the same midgut. We examined whether differences in fluorescence intensity correlated with parasite localization and time post infection in two mosquito species. To compare different experimental conditions, we normalized fluorescence intensity of each parasite based on the highest and lowest intensity of parasites in each image. We found only modest overall differences in mean fluorescence intensities at different positions (basal lamina, cellular layer, blood meal) over time and between the two species (*S7*-S9 Fig, Tables S8-S9). Furthermore, we observed the parasites with very low levels of fluorescence that appeared as a black hole on the background of the midgut cells expressing GFP reporter in *Ag* mosquitoes (Fig 5a) that expressed GFP uniformly in all midgut cells. In contrast, irregular patterns of GFP expression in the midgut were reported for *As* [18] (S1 Fig). Hence, fluorescence-negative parasites were only examined in *Ag* where parasites were clearly identified as black shapes on fluorescent background.

**Figure 5.**
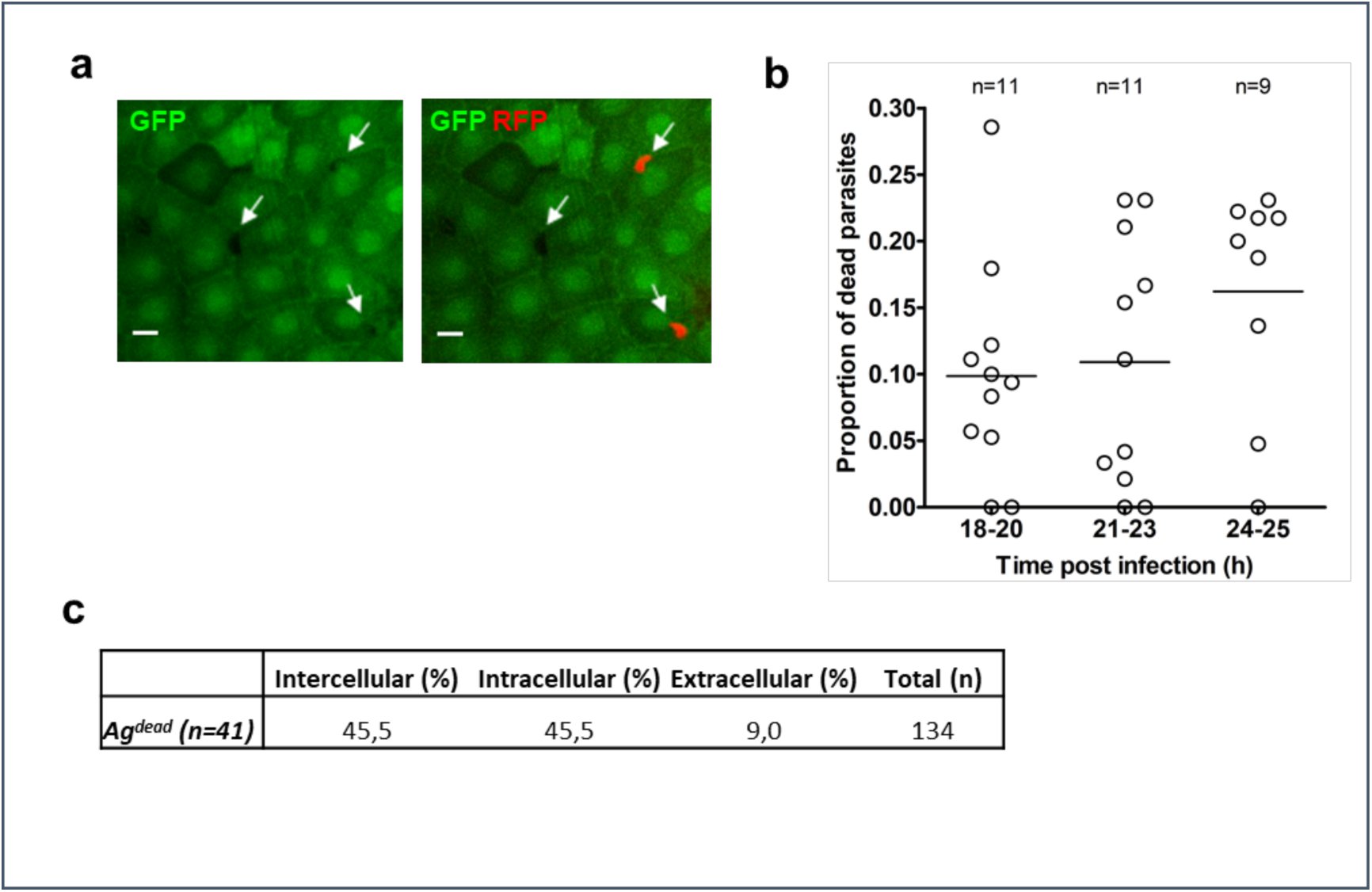
Quantification of dead parasites in *A. gambiae*. **a**. Detection of dead parasites within the cellular layer. Due to uniform GFP expression with the midgut cells of the *dmAct5C::GFP* line of *A. gambiae*, dead parasites that no longer express RFP could be distinguished in the midgut by their negative signal and a characteristic shape. Shown is a single z-section (scale bar - 20 µm) containing two live RFP-expressing parasites and one dead parasite, indicated by arrows. **b.** The proportion of dead parasites at different time points after *Ag* infection. Midguts (n) that contained at least 10 parasites were used for analyses. Each dot represents a single midgut. **c**. Distribution of dead parasites within the cellular layer. The table gives the number of parasites at each position at all time points. The number (n) is the number of midguts analyzed. Total (n) is the number of all analyzed parasites.

We considered the parasites that lost their fluorescence dying or dead [11,25]. On average, 10-15% of all recognized parasites had no fluorescence and were classified as dead (Fig 5b). Differences in distribution were observed for live and dead parasites within the cellular layer. More dead parasites were found to be located extracellular or intercellular (compare Fig 5c and Fig 4d). This observation points to more efficient parasite killing of extracellular parasites. Interestingly, we hardly detected any dead parasites in *Ag*^*TEP1KD*^ mosquitoes, suggesting that TEP1 may be involved in killing of parasites within the cellular layer.

### Cell damage caused by parasite passage

Midgut regeneration is a natural process of epithelia renovation after a blood feeding, whether infective or not [26]. Blood meal generates a stressful environment as it contains bacteria, reactive oxygen species and digestive enzymes that may cause damage to the midgut cells. It has been previously suggested that invaded midgut cells die after invasion and are expelled into the midgut lumen [27] resulting in accumulation of hundreds of cells in highly infected midguts. However, we only once observed GFP positive midgut cells in the midgut lumen. This result indicates that either upon expel dead midgut cells rapidly lose their GFP fluorescence, or that only few midgut cells are expelled after invasion. To resolve these conjectures, we investigated the integrity of the cell layer using high molecular weight Texas-Red conjugated dextran which is trapped inside damaged cells [28]. In these experiments, the fluorescent dextran was delivered into the midgut by blood feeding mosquitoes on mice injected intravenously with fluorescent dextran several minutes before mosquito feeding. We detected dextran filled cells (Fig 6a), calculated their position (Fig 6b, S11 Fig) and measured the distance to the nearest parasite (Fig 6c). The majority of dextran filled cells (70%) that contained a parasite in *As* were predominantly detected in the cellular layer. In contrast in *Ag*, dextran filled cells were observed both in the cellular layer and in the midgut lumen (Fig 6b). As many as 30% of dextran filled cells in *Ag*, were found in the midgut lumen. Out of these, 50% contained a parasite (S10 Table). In contrast, we found only one (5% of total) dextran filled cell the midgut lumen of *As* mosquitoes.

**Figure 6.**
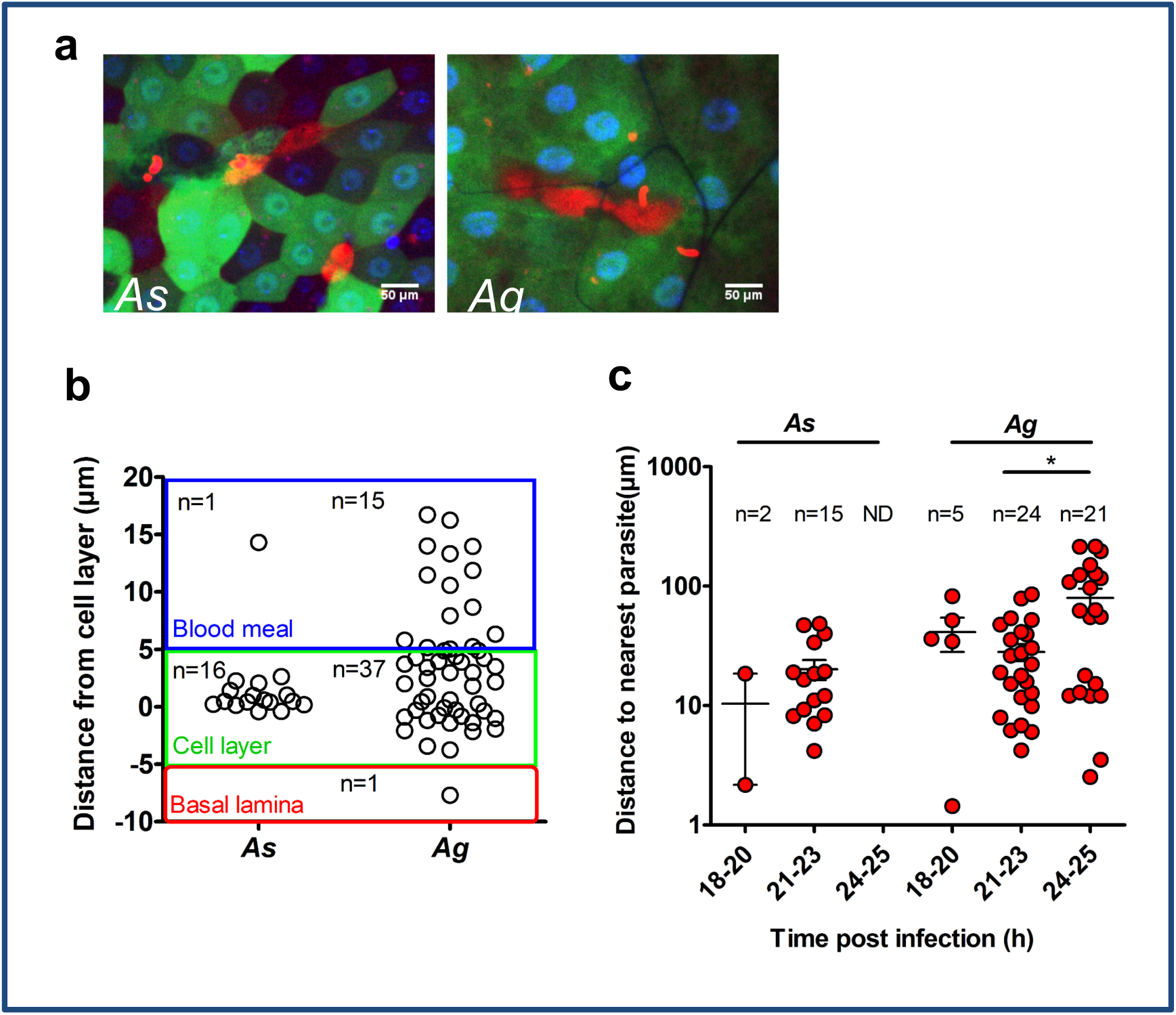
Quantification of damaged cells. **a**. Detection of dextran-positive cells in the midguts of *As* and *Ag* mosquitoes. Shown are single z-sections of GFP-expressing dissected midguts. Mosquitoes were fed on mice injected with Texas-Red conjugated dextran. Dextran-filled cells appeared red (scale bar - 50 µm). **b**. Positions of dextran-filled cells in the midgut layers of *As* and *Ag*. Each dot represents a single dextran-positive cell. The graph depicts positions of the dextran-positive cells within the midgut layers. Each midgut layer is color coded: Blood meal (blue), cellular layer (green) and basal lamina (red). The number of dextran filled cells (n) at each position is indicated. **c.** Distances of dextran-filled cells to the nearest parasite at different time points after infection of *Ag* and *As*. The number of dextran filled cells analyzed (n) is shown. Statistical analysis was performed by a Mann-Whitney non-parametric t-test.

Interestingly in *Ag*, the distance between the dextran-positive cell and the nearest parasite significantly increased at 24-25 hpi compared to the earlier time intervals (Fig 6c). Furthermore, no dextran-positive cells were found in *As* at the late time interval after infection (24-25 hpi). Taken together, these results suggest that in *Ag* mosquitoes, damaged cells are readily extruded into the midgut lumen with or without the parasites. It is important to note that while some dextran filled cells contained a parasite, most midgut cells that we observed to host a parasite were dextran-negative, indicating that ookinete invasion damaged and killed only a small proportion of midgut cells. Interestingly, in both *As* and *Ag*, we never observed more than one parasite in a non-damaged midgut cell, indicating that parasites refrain from entering an invaded cell. Here we also observed chains of several connected dextran-positive cells, indicating that ookinetes can traverse several neighboring cells before exiting on the basal side of the cellular layer. In conclusion, our results led us to suggest that the route of ookinete invasion for the same parasite is species-specific and shaped by midgut tissue morphology, physiology, damage and immune responses. Future studies should examine how invasion strategies of the human malaria *P. falciparum* parasites are affected by diverse vector species.

## Conclusions

By combining live imaging techniques with quantitative bioimage analysis workflow, we uncovered differences in ookinete invasion strategies in two related mosquito species. We showed that in both species, the “pioneer” parasites that first reach the basal side of the midgut were rapidly eliminated by the mosquito immune system, and that colonization of the mosquito midgut was initiated at later stages of the infection. High throughput image data analyses of two *Anopheles* species revealed important differences in parasite invasion routes. We showed that the average ookinete speed in the cellular layer is lower in *As* compared to *Ag* mosquitoes. Moreover, *As* midguts contained more intercellular parasites and displayed higher numbers of damaged parasite-harboring cells. These results indicate that faster ookinete speeds and preference for intracellular route may impede parasite survival during invasion in *Ag*, the mosquito species which is more resistant to *P. berghei* infection. The reported here combination of live imaging and automated image analysis is highly adaptable and can be extended to functional analyses of gene knockdowns, mutations, and drug treatments. Moreover, the image data base and image analysis tools generated by this study offer a powerful tool for studying *Plasmodium* motility in *Anopheles* mosquitoes.

## Materials and methods

### Mosquito rearing

Transgenic *Anopheles stephensi* mosquitoes expressing GFP under the midgut-specific G12 promoter (*pG12::EGFP transgenic line* [18]) and *Anopheles gambiae* expressing GFP under the *Drosophila Acti5c* promoter (*dmActin5c::dsx-eGFP*) line [19]) were reared in the lab as previously described [29]. Briefly, mosquitoes were maintained in standard conditions (28°C, 75– 80% humidity, 12-hr/12-hr light/dark cycle). Larvae were raised in deionized water and fed finely ground TetraMin fish food. Adults were fed on 10% sucrose *ad libitum* and females were blood fed on anaesthetized mice. To obtain *Ag* mosquitoes that do not express *TEP1*, the dominant *TEP1* knockdown *Ag*^*TEP1KD*^ transgenic line [30] was crossed to *dmActin5c::dsx-*e*GFP* mosquitoes. The F1 progeny had reduced TEP1 levels while expressing GFP in the midgut [30].

### *P. berghei* infections

For infections, mosquitoes were blood fed on *P. berghei* infected mice as previously described [31]. *P. berghei* pyrimethamine resistant strain (RMgm 296) constitutively expressed RFP [20]. For the visualization of damaged mosquito cells, mice were injected in the tail vein with 0.1 ml of 5% dextran (3,000 kDa Texas Red conjugated, Invitrogen) diluted in PBS 10 min prior to blood feeding. Mosquitoes were blood fed for 20 min on anesthetized mice and dissected between 18-24 h after blood feeding, as indicated in each experiment.

### Confocal microscopy

Immediately prior to visualization, infected mosquitoes were dissected on ice in PBS buffer supplemented with 0.02% DAPI (Thermo Fisher, 4′,6-diamidino-2-phenylindole, 5 mg/mL), and with 0.2% tricaine (Sigma), 0.02% tetramisole (Sigma) to prevent midgut contraction during image acquisition. Blood-filled midguts were placed on 35 mm plastic dishes with glass bottom (Nunc, ThermoFisher). Dishes were mounted on inverted DMI6000 Leica Microscope, equipped with a Nipkow Disk confocal module (Andor Revolution), 20X objective. For time-lapse experiments, samples were visualized for up to two hours at 1 min intervals. The number of 1 µm-stacks, annotated for each image, ranged between 24 and 95 depending on tissue thickness. We noticed that *As* midguts were rigid and sturdy, allowing for longer live imaging. *Ag* midguts were more fragile and tended to move and tear during image acquisition. We were able to collect live imaging data from 16 *As* (S1 Table) and 5 *Ag* midguts (S2 Table). We were not able to follow parasites in mosquitoes lacking the immune protein TEP1 due to high midgut fragility of *Ag*^*TEP1KD*^ midguts.

### Image analyses

All images were uploaded to a database where they were annotated according to mosquito species and other experimental conditions. Images were subjected to bulk analysis as well as manual verification. The annotated image database is accessible to JAVA programming using the Strand Avadis IManage data management software. All data (images and extracted data as text files) are available on cid.curie.fr, Project “Malaria parasite invasion in the mosquito tissues” at https://cid.curie.fr/iManage/standard/login.html. The META data is managed using OpenImadis https://strandls.github.io/openimadis/. Companion scripts are available here: https://github.com/PerrineGilloteaux/MalariaParasiteinMosquito.

The api documentation is available under API tab https://cid.curie.fr/iManage/api/client/. The api client jar is available at https://cid.curie.fr/iManage/standard/downloads.html.

Companion scripts include segmentation of parasites, nuclei and quantification of intensities corrected by background was performed using a set of ImageJ Plugins in Java. Analysis of the position of parasites relative to cell layer and statistics are performed with MATLAB.

The data set was collected from 110 experiments including a total of 2,557 parasites (*As* - 45 midguts, 1,068 parasites; *Ag* - 34 midguts, 796 parasites; and *Ag*^*Tep1KD*^ – 31 midguts, 693 parasites). There was no bias in the number of parasites per midgut across different time points and mosquito species (S2 Fig). Furthermore, we found that infection levels (low, intermediate, or high) had some effect on the result of parasite distribution, specifically in at low infection levels (S3 Fig). Consequently, we used for our analysis only images that contained at least 10 parasites per image.

### Ethics statement

The animal work described in this study received agreement #E67-482-2 from the veterinary services of the region Bas-Rhin, France (Direction départementale des services vétérinaires).

## Acknowledgments

GV, JŠ, JS and EAL thank M.E. Moritz and C. Kappler for help with the mosquito colony and parasite cultures; and E. Marois for scientific discussions and support. JeS and PPG acknowledge the Structure fédérative de recherche santé François-Bonamy and the SERPICO team, are members of the national infrastructure “France BioImaging”. Authors thank A. Volohonsky for graphic expertise.

## Author contributions

Conceived and designed the experiments: GV, EAL. Performed the experiments: GV, JŠ, JuS. Analyzed the data: GV, PPG, JeS, EAL. Contributed reagents/materials/analysis tools: PPG, JeS. Wrote the paper: GV, EAL.

## Notes

#### Summary of Updates

The correct name spelling and ORCID# of Jitka Stafkova.

https://cid.curie.fr/iManage/standard/login.html

https://strandls.github.io/openimadis/

https://github.com/PerrineGilloteaux/MalariaParasiteinMosquito

https://cid.curie.fr/iManage/api/client/

https://cid.curie.fr/iManage/standard/downloads.html

