## Supplemental materials for "Kinetics of *Plasmodium midgut* invasion in Anopheles mosquitoes"

**Short Title: Live imaging of *Plasmodium* invasion of mosquito midguts**

Gloria Volohonsky[1]*, Perrine Paul-Gilloteaux[2,4], Jitka Štáfková[1,5], Julien Soichot[1,6], Jean Salamero[2], Elena Levashina[1,3]*

#### Supplementary Figures

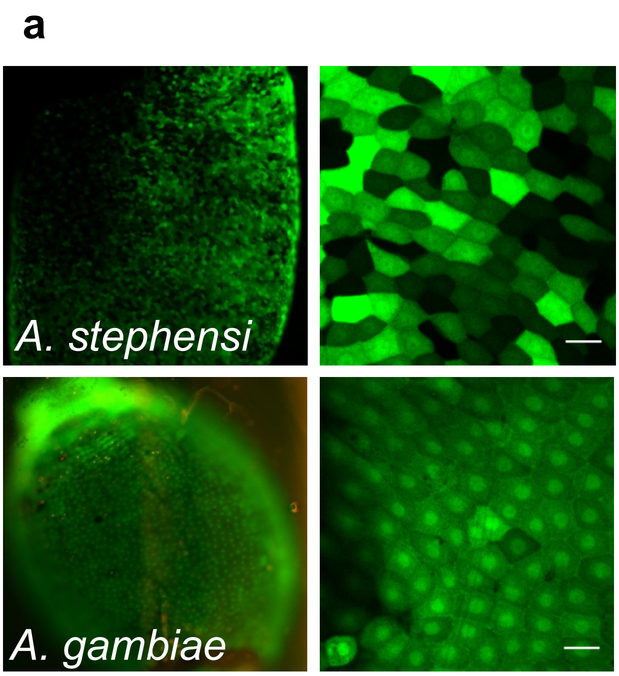


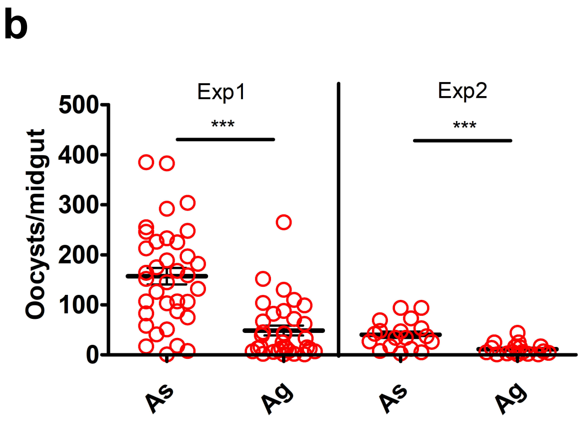


| | Exp 1 | | Exp 2 | |
|---|---|---|---|---|
| Parasitemia (%) | 5 | | 3 | |
| | ***As*** | ***Ag*** | ***As*** | ***Ag*** |
| Prevalence (%) | 100 | 92 | 100 | 68 |
| Median oocysts/midgut | 156 | 25 | 35 | 6 |
| *p value* | <0.0001 | | 0.0005 | |

**Figure S1. *P. berghei* infection intensities in the transgenic *A. stephensi* and *A. gambiae* mosquitoes expressing GFP in the midgut cells. a.** GFP fluorescence in the midgut cells of *A. stephensi pG12::EGFP* line (upper) and *A. gambiae dmActin5c::dsx-eGFP* line (lower) 24 h after blood feeding. Enlarged are representative 20-fold magnification images showing GFP fluorescence in enterocytes (scale bar - 50 μm). **b.** *P. berghei* infection intensities in *A. stephensi* and *A. gambiae*. Oocysts were counted in dissected midguts 7 days post infection. The results of two independent experiments are shown. Prevalence indicates the percentage of infected mosquito midguts in each experiment. Horizontal lines depict median number of oocysts per midgut. Statistical differences between infections of *As* and *Ag* were evaluated by a nonparametric *t* test, *** indicate $p < 0.0005$.

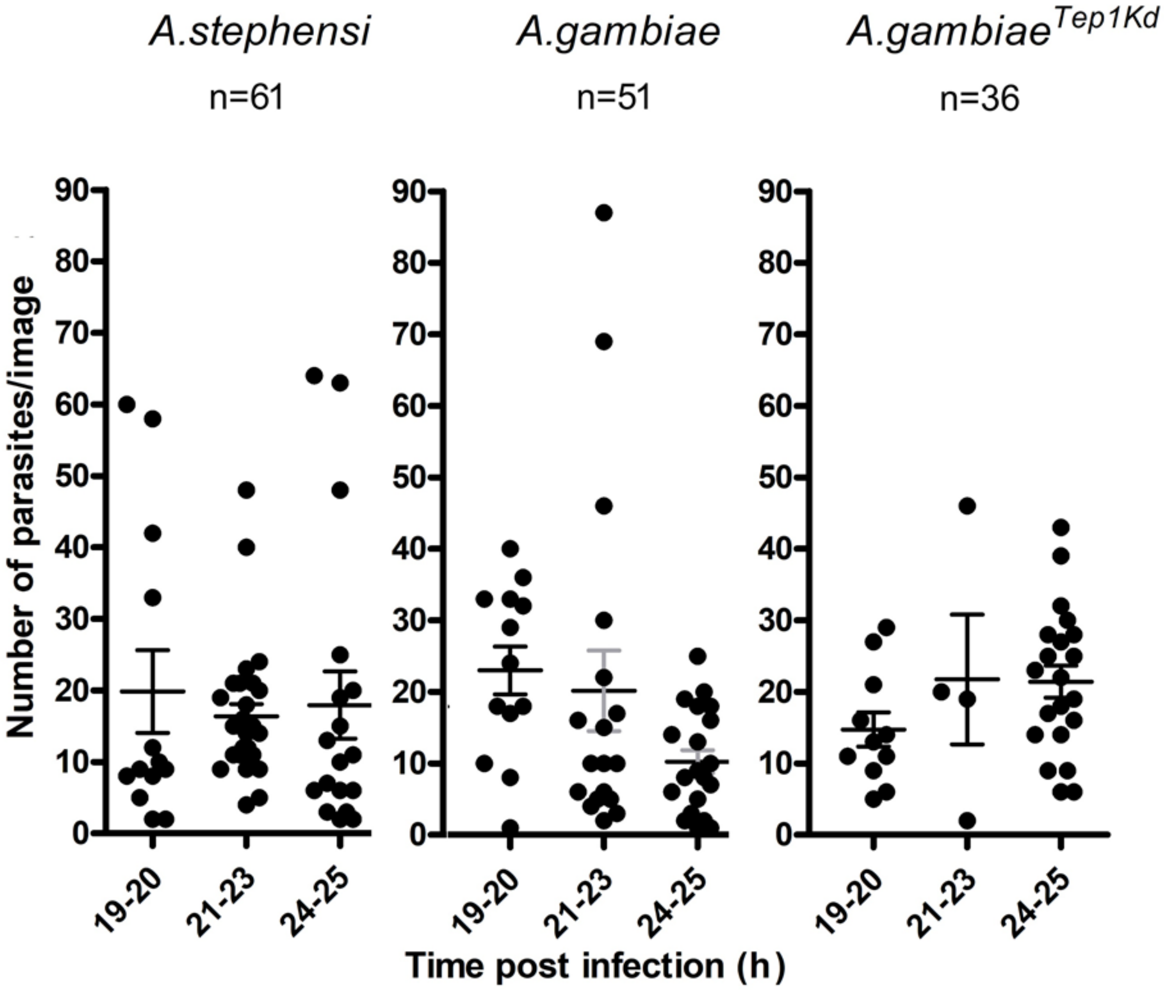


**Figure S2**. Parasite numbers used for analyses in *A. stephensi*, *A. gambiae*, and *A. gambiae* depleted for TEP1 (*A. gambiae*$^{TEP1KD}$). Each dot represents a single midgut. Similar numbers of parasites were analyzed in all mosquitoes at the indicated time points after infection (hpi). n= The number of midguts analyzed.

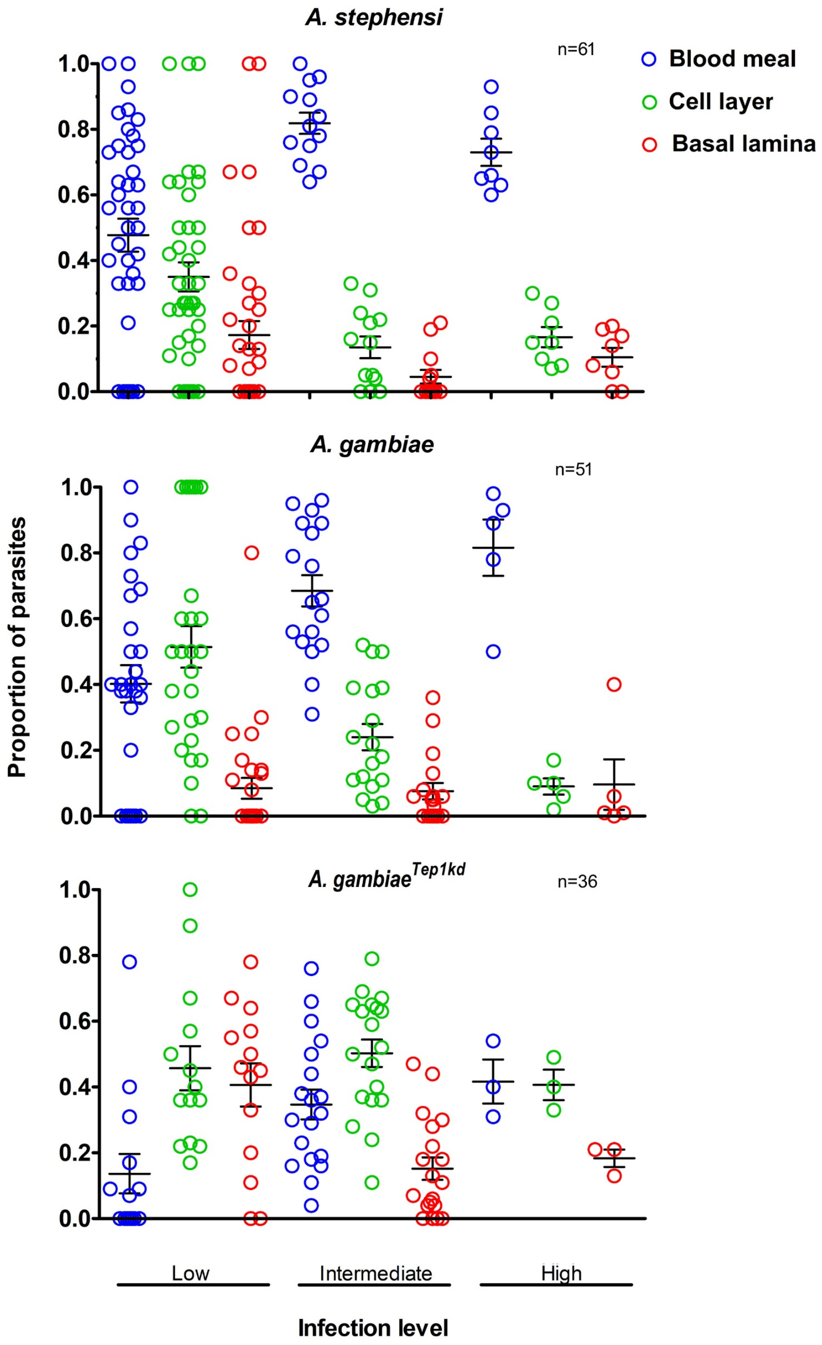


**Figure S3**. Parasite distribution in *A. stephensi*, *A. gambiae*, and *A. gambiae* depleted for TEP1 (*A. gambiae*$^{TEP1KD}$) at low, intermediate and high infection levels. Blood meal (blue), cellular layer (green) and basal lamina (red) in midguts with different infection levels. Low infection (up to 15 parasites), intermediate (16-35) and high (more than 35 parasites per image) are compared. Each dot represents the proportion of parasites at a given position in a single midgut. *n*= number of analyzed images.

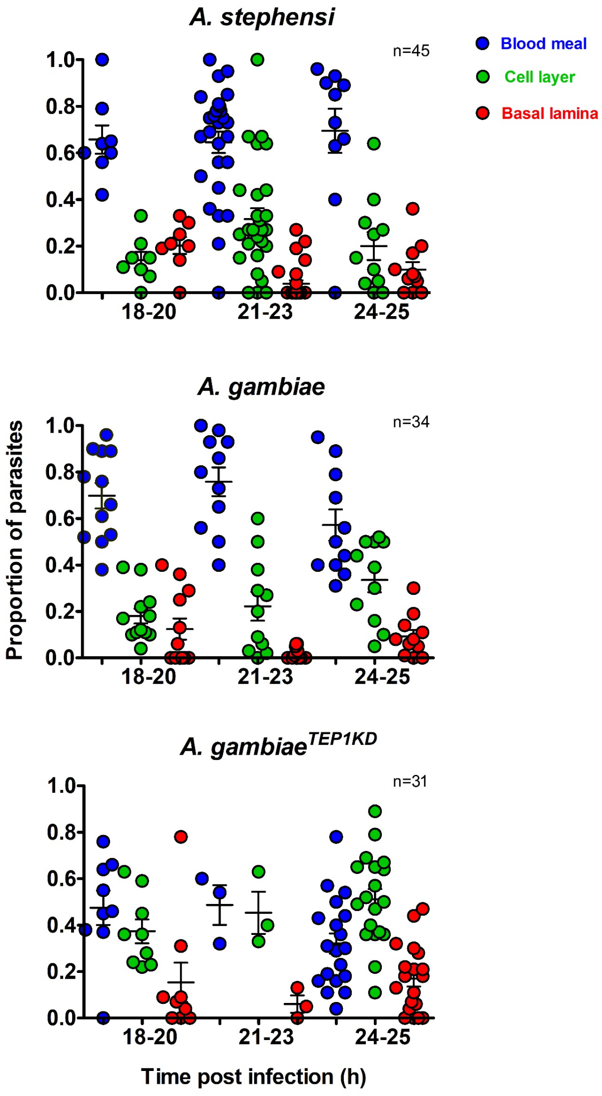


**Figure *S4*.** Time dependent parasite distribution in *A. stephensi*, *A. gambiae*, and *A. gambiae* depleted for TEP1 (*A. gambiae$^{TEP1KD}$*). Proportion of parasites found in blood meal (blue), cellar layer (green) and basal lamina (red) at the indicated time points after infection (hpi). *N* = number of analyzed images. Analyzed images containing at least ten parasites.

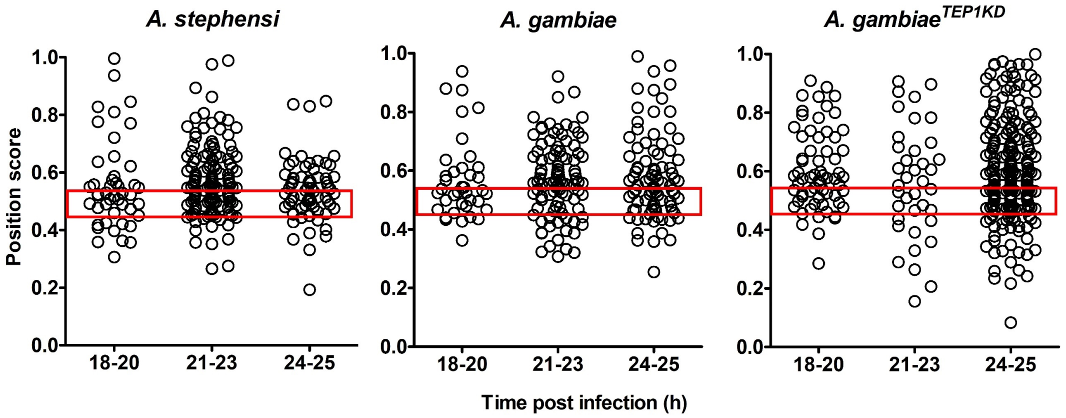


**Figure S5**. Parasite distribution within the cellular layer in *A. stephensi*, *A. gambiae*, and *A. gambiae* depleted for TEP1 (*A. gambiae*$^{TEP1KD}$). Scatter plots depict the score for each parasite at indicated times after infection. Parasites are considered extracellular when the score $s < 0.45$, intercellular for the score $0.45 < s < 0.55$ (red box) and intracellular if the score $s > 0.55$. n= number of parasites analyzed at each time interval.

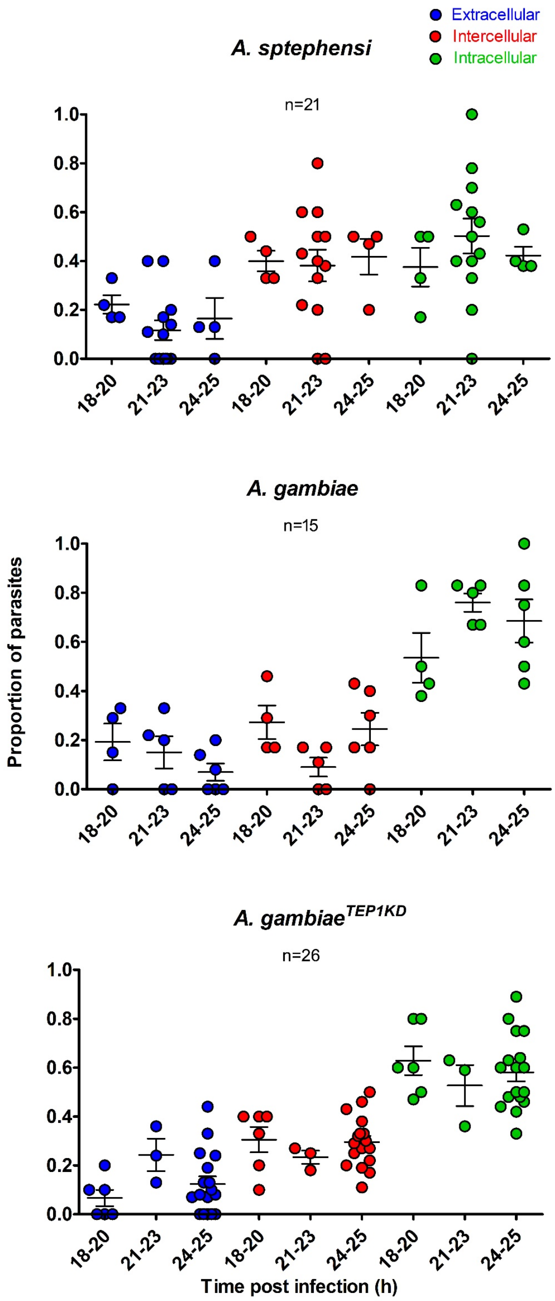


**Figure S6**. Parasite positions within the cellular layer in *A. stephensi*, *A. gambiae*, and *A. gambiae* depleted for TEP1 (*A. gambiae*$^{TEP1KD}$). Scatter plots depict the proportion of parasites at each position within the cell layer: extracellular (blue), intercellular (red) and intracellular (green) at different time intervals after infection. Each dot represents a single image which contained at least six parasites in the cellular layer. n= number of analyzed images.

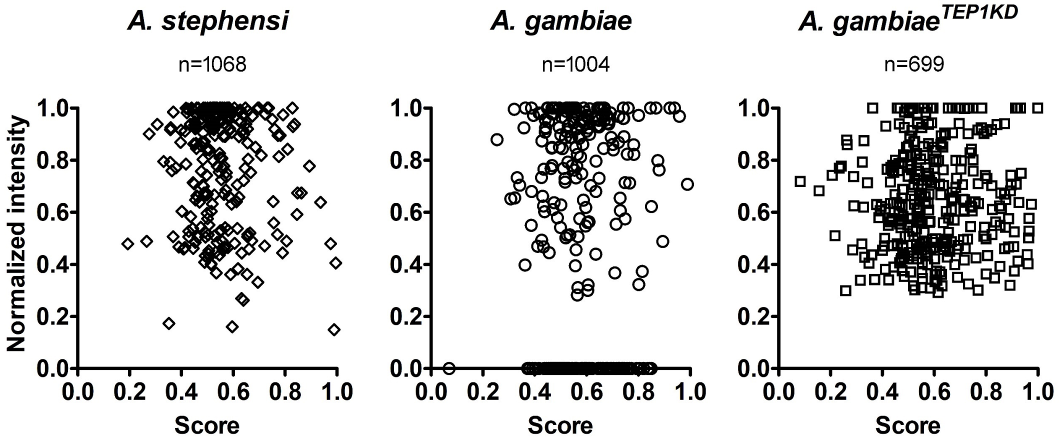


**Figure S7**. Intensity of parasite fluorescence versus position score in *A. stephensi*, *A. gambiae*, and *A. gambiae* depleted for TEP1 (*A. gambiae*$^{TEP1KD}$). Each dot represents a parasite. n= number of analyzed parasites. No correlation was found between the position of the parasite and the level of fluorescence intensity.

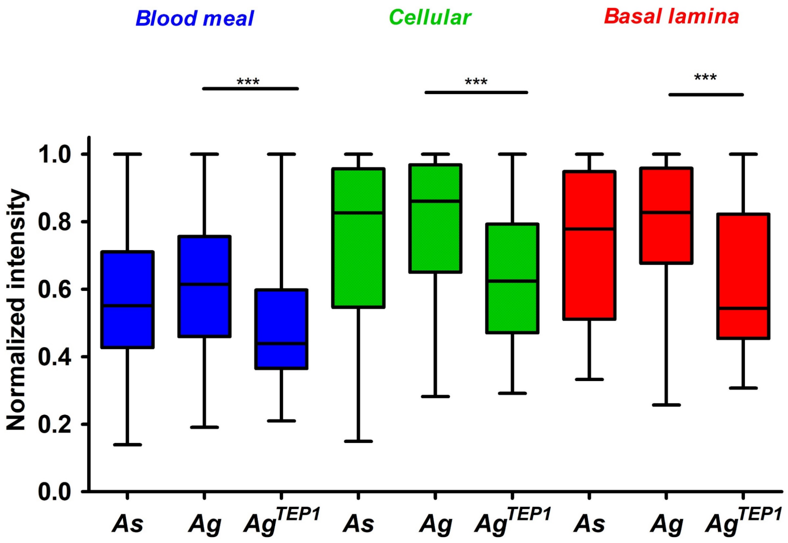


**Figure S8.** Intensity of parasite fluorescence in *A. stephensi* (*As*), *A. gambiae* (*Ag*), and *A. gambiae* depleted for TEP1 ($Ag^{TEP1KD}$). Bar graph depicts the distribution of parasite fluorescence intensity at different positions: blood meal (blue), cellular layer (green) and basal lamina (red). Parasites from all time points were pooled to calculate the average normalized intensity. Parasite intensity is normalized for each image, intensity ranges between 0.0 and 1.0, where 1.0 is the maximum intensity observed. Statistical significance of differences within each group was tested by one-way ANOVA, $p < 0.0001$.

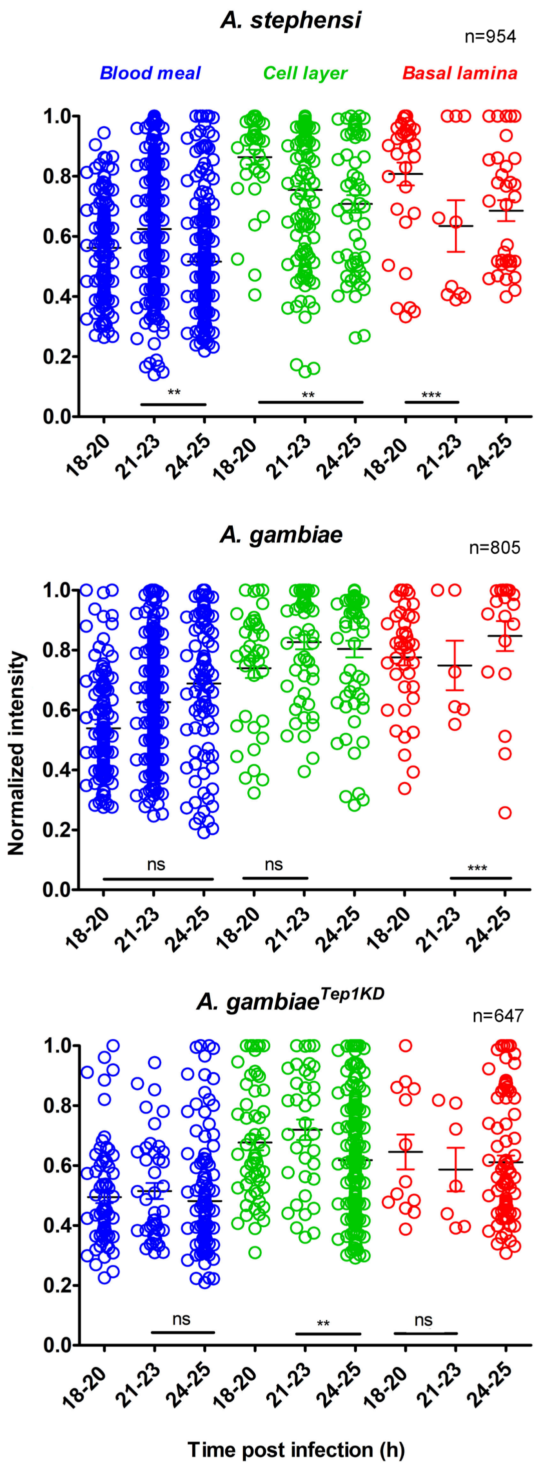


**Figure S9.** Intensity of parasite fluorescence in *A. stephensi* (*As*), *A. gambiae* (*Ag*), and *A. gambiae* depleted for TEP1 ($Ag^{TEP1KD}$) at different times after infection and at different positions: blood meal (blue), cellular layer (green) and basal lamina (red). Each circle represents a parasite. n= number of analyzed parasites. Statistical analysis was performed using non-parametrical Mann Whitney test. Only images with more than 10 parasites were analyzed.

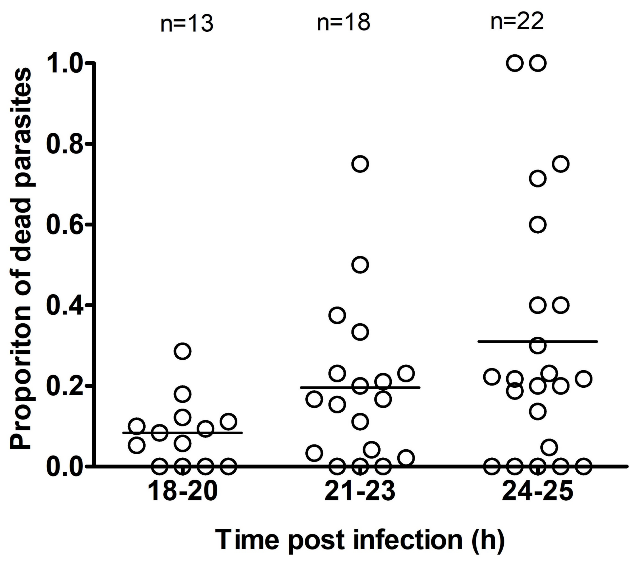


**Figure S10.** Quantification of dead parasites in *A. gambiae*. The proportion of parasites that were considered dead in each image at indicated time intervals after infection. Each dot represents an image. n= the number of analyzed images.

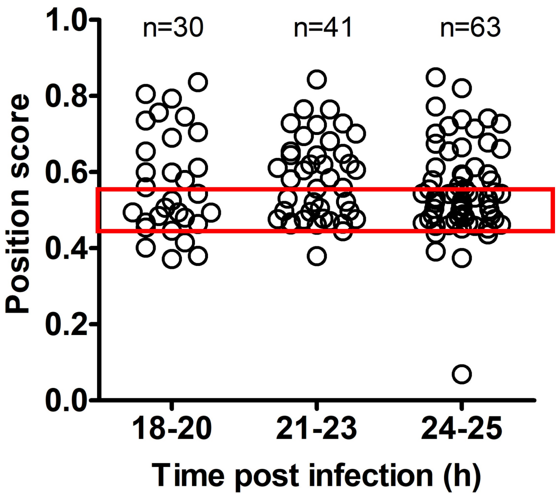


**Figure S11.** Distribution of dead parasites within the cellular layer in *A. gambiae*. Scatter plots depict the score for each parasite at indicated times after infection. Parasites are considered extracellular when the score $s < 0.45$, intercellular for the score $0.45 < s < 0.55$ (red box) and intracellular if the score $s > 0.55$. n = number of parasites analyzed at each time interval.

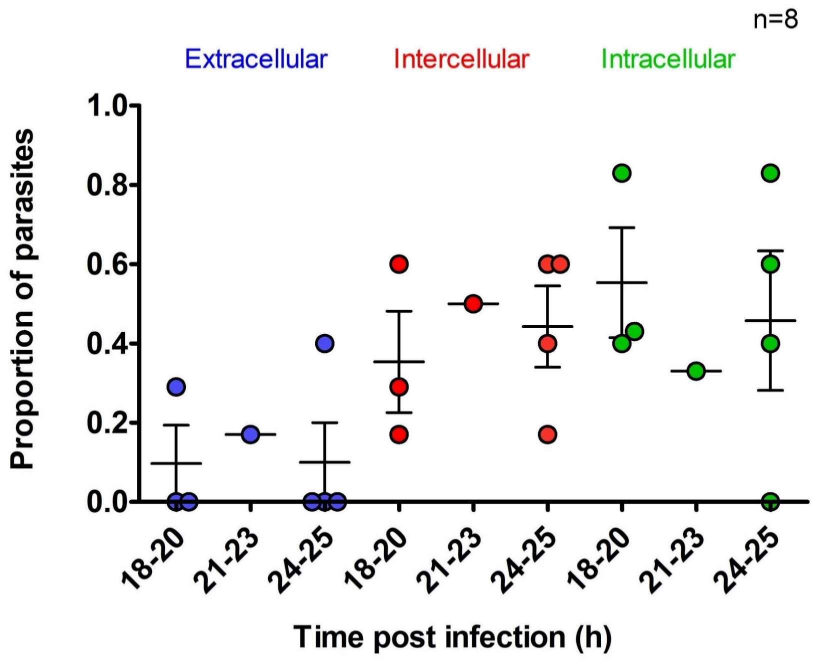


**Figure S12**. Position of dead parasites in *A. gambiae* within the cellular layer. Scatter plots depict the proportion of parasites at each position within the cell layer: extracellular (blue), intercellular (red) and intracellular (green) at different time intervals after infection. Each dot represents a single image (n = 8).

### Supplementary Tables

**Table S1.** Time-lapse records of ookinete invasion of *A. stephensi* midguts in the presence or absence of dextran, the marker of cell membrane integrity. Guid - the unique record number in the database, tpi - hours post infection, duration - the total time of the time-lapse measurement.

| Guid | Dextran | hpi (h) | duration(min) | Feature | link |
|---|---|---|---|---|---|
| 2114 | no | 19 | 22 | **Z-project** | https://youtu.be/1_KEdIZWcqE |
| 2115 | no | 20 | 120 | **Z-project** | https://youtu.be/wkKHlYjaFwI |
| 1680 | no | 20 | 89 | **Z-project** | https://youtu.be/jfSG_acwGB0 |
| 1774 | no | 20 | 85 | **Z-project** | https://youtu.be/ROu7OrQ2w6I |
| | | | | **overview** | https://youtu.be/xVSNqvvbvDw |
| | | | | sideview | https://youtu.be/Up-RM_cicGo |
| | | | | sideview | https://youtu.be/w7TyNLBEeWA |
| | | | | sideview | https://youtu.be/a1HRxJlrzQU |
| | | | | zoom in | https://youtu.be/GDONJcZcCzA |
| 2107 | yes | 20 | 92 | **Z-project** | https://youtu.be/1vVwNaXOGYs |
| 0 | yes | 22 | 120 | **Z-project** | https://youtu.be/uRYpQGjh_vA |
| 2112 | no | 22 | 112 | **Z-project** | https://youtu.be/Hck5ksxpRvs |
| | | | | **overview** | https://youtu.be/SDpQ8KHQ2_M |
| | | | | zoom in | https://youtu.be/iaDC58NetIk |
| | | | | zoon in, rotate | https://youtu.be/DNb44lSTn8M |
| 1775 | no | 22.5 | 42 | **Z-project** | https://youtu.be/IsNrlOz1t-w |
| | | | | **overview** | https://youtu.be/C8mHzXEqIXo |
| | | | | zoom in | https://youtu.be/TJ8e3HhbxRk |
| | | | | zoom in | https://youtu.be/xsiwNR9zAPM |
| | | | | zoom in | https://youtu.be/1NBKAUeoBAo |
| 2119 | no | 22.5 | 95 | **Z-project** | https://youtu.be/ethg8Awj5FU |
| | | | | Overview + sideview | https://youtu.be/yGCnEb8UvJs |
| | | | | zoom in | https://youtu.be/FQy9jCkCalk |
| 1628 | no | 22.5 | 115 | **Z-project** | https://youtu.be/VpoFhhtUze0 |
| 1679 | no | 24 | 54 | **Z-project** | https://youtu.be/voohki6Qt0o |
| 2111 | | | | **Z-project** | https://youtu.be/52ZajjJXpsc |
| | | | | overview | https://youtu.be/Nl2Wk5zjnNw |
| | | | | sideview | https://youtu.be/bd-qGHj4KJ0 |
| 1777 | no | 24 | 58 | **Z-project** | https://youtu.be/NdMDozl9w5M |
| | | | | zoom in | https://youtu.be/UHNmziUt7iM |
| 1615 | no | 24.5 | 33 | **Z-project** | https://youtu.be/QlQrrtWgp9Q |
| | | | | sideview | https://youtu.be/r6Wcr7D10oo |
| | | | | overview | https://youtu.be/X1E31lS5gUA |
| 2113 | no | 24.5 | 49 | **Z-project** | https://youtu.be/DXH4xMxqCHc |
| 0 | no | 26 | 120 | **Z-project** | https://youtu.be/Z-zhEGavE34 |
| | | | | overview | https://youtu.be/oGMtzqZduJ8 |
| | | | | sideview | https://youtu.be/iO5OP0srrVw |

**Table S2.** Time-lapse records of ookinete invasion of *A. gambiae* midguts in the presence of dextran, the marker of cell membrane integrity. Guid - the unique record number in the database, hpi - hours post infection, duration - the total time of the time-lapse measurement.

| Guid | Dextran | hpi | Duration min) | Feature | Link |
|---|---|---|---|---|---|
| 1796 | Yes | 19 | 50 | **Z-project** | *https://youtu.be/RuxaIPgHywo* |
| 1621 | Yes | 20 | 0 | rotation of a single image | *https://youtu.be/ie-9RJ7t_pg* |
| 2110 | Yes | 20.5 | 77 | **overview** | *https://youtu.be/Zffuz1oevKk* |
| 1622 | Yes | 21.5 | 48 | **Z-project** | *https://youtu.be/4I5wZgSPOzA* |
| 2109 | Yes | 22 | 31 | **Z-project** | *https://youtu.be/m3VRyDDcxl4* |
| 1624 | Yes | 22.5 | 20 | **Z-project** | *https://youtu.be/bNN1Bu6-fOA* |
| 2108 | Yes | 24.5 | 63 | **Z-project** | *https://youtu.be/UlK_dlvEHts* |
| | | | | diagonal view | *https://youtu.be/KRtBQAXlR3A* |
| | | | | side view | *https://youtu.be/qnBLe2kpWGM* |

**Table S3.** Kruskal-Wallis test of differences in ookinete localization between *A. gambiae* (*Ag*) and *A. gambiae* with silenced *TEP1* ($Ag^{TEP1KD}$) at the indicated time points post infection (hpi).

| Ookinete distribution | Kruskal Wallis test | | |
|---|---|---|---|
| | ***Ag*** | $\boldsymbol{Ag^{TEP1KD}}$ | *p-value* |
| **18-20 hpi** | | | |
| Blood meal | ns | ns | 0.2186 |
| Cellular layer | < $Ag^{TEP1KD}$ | > *Ag* | 0.0418 |
| Basal lamina | ns | ns | 0.8842 |
| **21-23 hpi** | | | |
| Blood meal | > $Ag^{TEP1KD}$ | <*Ag* | 0.064 |
| Cellular layer | < $Ag^{TEP1KD}$ | > *Ag* | 0.0826 |
| Basal lamina | ns | ns | 0.4561 |
| **22-25 hpi** | | | |
| Blood meal | > $Ag^{TEP1KD}$ | < *Ag* | 0.0049 |
| Cellular layer | < $Ag^{TEP1KD}$ | > *Ag* | 0.0217 |
| Basal lamina | ns | ns | 0.5752 |

**Table S4**
Kruskal-Wallis test of differences in parasite localization in *A. stephensi* (*As*), *A. gambiae* (*Ag*) and *A. gambiae* silenced for *TEP1* (*Ag*$^{TEP1KD}$) between the indicated time points after infection (hpi).

| Ookinete position | Kruskal Wallis test | | | |
|---|---|---|---|---|
| | **18-20 hpi** | **21-23 hpi** | **24-25 hpi** | *p-value* |
| ***As*** | | | | |
| Blood meal | ns | ns | ns | 0.5852 |
| Cellular layer | ns | ns | ns | 0.1077 |
| Basal lamina | > 21-23 hpi | < 18-20 hpi<br>< 24-25 hpi | >21-23 hpi | 0.0005 |
| ***Ag*** | | | | |
| Blood meal | ns | >24-25 hpi | < 21-23 hpi | 0.0369 |
| Cellular layer | ns | < 24-25 hpi | > 21-23 hpi | 0.0634 |
| Basal lamina | ns | < 24-25 hpi | > 21-23 hpi | 0.0415 |
| ***Ag*$^{TEP1KD}$** | | | | |
| Blood meal | > 24-25 hpi | ns | < 18-20 hpi | 0.044 |
| Cellular layer | < 24-25 hpi | ns | > 18-20 hpi | 0.0499 |
| Basal lamina | ns | ns | ns | 0.6019 |

**Table S5.** Kruskal-Wallis test of differences in parasite localization between *A. stephensi* (*As*), *A. gambiae* (*Ag*) and *A. gambiae* silenced for *TEP1* (*Ag*$^{TEP1KD}$) at the indicated time points after infection (hpi).

| Ookinete position | | | Kruskal Wallis test | |
|---|---|---|---|---|
| | ***As*** | ***Ag*** | ***Ag***$^{TEP1KD}$ | p-value |
| **18-20 hpi** | | | | |
| Blood meal | ns | ns | ns | 0.3457 |
| Cellular layer | ns | ns | ns | 0.1291 |
| Basal lamina | ns | ns | ns | 0.4012 |
| **21-23 hpi** | | | | |
| Blood meal | ns | >*Ag*$^{Tep1KD}$ | <*Ag* | 0.0833 |
| Cellular layer | ns | ns | ns | 0.125 |
| Basal lamina | ns | ns | ns | 0.3879 |
| **24-25 hpi** | | | | |
| Blood meal | > *Ag*$^{TEP1KD}$ | > *Ag*$^{TEP1KD}$ | < *Ag*,< *As* | 0.0033 |
| Cellular layer | < *Ag*$^{TEP1KD}$ | < *Ag*$^{TEP1KD}$ | > *Ag*,> *As* | 0.0001 |
| Basal lamina | ns | ns | ns | 0.7941 |

**Table S6.** Kruskal-Wallis test of differences in parasite fluorescence intensities in *A. stephensi* (*As*), *A. gambiae* (*Ag*) and *A. gambiae* silenced for *TEP1* ($Ag^{TEP1KD}$) between the indicated time points after infection (hpi).

| Ookinete fluorescence intensity | | | | Kruskal Wallis test |
|---|---|---|---|---|
| | **18-20 hpi** | **21-23 hpi** | **24-25 hpi** | *p-value* |
| ***As*** | | | | |
| Blood meal | >24-25 hpi<br><21-23 hpi | >18-20 hpi<br>>24-25 hpi | <18-20 hpi<br><21-23 hpi | |
| Cellular layer | >24-25 hpi | ns | <18-20 hpi | 0,01 |
| Basal lamina | >24-25 hpi | ns | <18-20 hpi | 0,037 |
| ***Ag*** | | | | |
| Blood meal | <21-23 hpi<br>< 24-25 hpi | >18-20 hpi | >18-20 hpi | 1,29E-08 |
| Cellular layer | <21-23 hpi | >18-20 hpi | ns | 0,045 |
| Basal lamina | ns | ns | ns | 0,1171 |
| $\boldsymbol{Ag^{TEP1KD}}$ | | | | |
| Blood meal | ns | ns | ns | 0,313 |
| Cellular layer | ns | >24-25 hpi | <21-23 hpi | 0,008 |
| Basal lamina | ns | ns | ns | 0,648 |

**Table S7.** Kruskal-Wallis test of differences in parasite fluorescence intensities between *A. stephensi* (*As*), *A. gambiae* (*Ag*) and *A. gambiae* with silenced *TEP1* ($Ag^{TEP1KD}$) at all time points.

| Ookinete intensity | | | | Kruskal Wallis test |
|---|---|---|---|---|
| | ***As*** | ***Ag*** | $\boldsymbol{Ag^{TEP1KD}}$ | *p-value* |
| Blood meal | > $Ag^{TEP1KD}$< *Ag* | >$Ag^{TEP1KD}$> *As* | < *Ag*, *As* | 2,00E-17 |
| Cellular layer | > $Ag^{TEP1KD}$ | > $Ag^{TEP1KD}$ | < *Ag*, *As* | 1,00E-15 |
| Basal lamina | > $Ag^{TEP1KD}$ | > $Ag^{TEP1KD}$ | < *Ag*, *As* | 2,48E-06 |

**Table S8.** Kruskal-Wallis test of differences in parasite fluorescence intensities between *A. stephensi* (*As*), *A. gambiae* (*Ag*) and *A. gambiae* silenced for *TEP1* (*$Ag^{TEP1KD}$*) at the indicated time points after infection (hpi).

| Ookinete fluorescence intensity | | | | Kruskal Wallis test |
|---|---|---|---|---|
| | ***As*** | ***Ag*** | ***$Ag^{TEP1KD}$*** | *p-value* |
| **18-20 hpi** | | | | |
| Blood meal | > *$Ag^{TEP1KD}$* | ns | *$Ag^{TEP1KD}$* < *As* | 0,00 |
| Cellular layer | > *$Ag^{TEP1KD}$* | ns | *$Ag^{TEP1KD}$* < *As* | 6,10E-05 |
| Basal lamina | > *$Ag^{TEP1KD}$* | ns | < *As* | 0,03 |
| **21-23 hpi** | | | | |
| Blood meal | > *$Ag^{TEP1KD}$* | > *$Ag^{TEP1KD}$* | < *Ag, As* | 5,00E-04 |
| Cellular layer | ns | > *$Ag^{TEP1KD}$* | < *Ag* | 0,02 |
| Basal lamina | ns | ns | ns | 0,62 |
| **24-25 hpi** | | | | |
| Blood meal | < *Ag* | > *$Ag^{TEP1KD}$*,> *As* | < *Ag* | 3,00E-10 |
| Cellular layer | > *$Ag^{TEP1KD}$* | > *$Ag^{TEP1KD}$* | < *Ag, As* | 6,00E-08 |
| Basal lamina | ns | > *$Ag^{TEP1KD}$* | < *Ag* | 2,60E-03 |

**Table S9**. Kruskal-Wallis analyses of parasite fluorescence intensities in *A. stephensi* (*As*), *A. gambiae* (*Ag*) and *A. gambiae* silenced for *TEP1* ($Ag^{TEP1KD}$), between the different positions blood meal (BM), cellular layer (CL) and basal lamina (BL) at the indicated time points after infection (hpi).

| Ookinete fluorescence intensity | | | | Kruskal Wallis test |
|---|---|---|---|---|
| | **Blood Meal** | **Cell Layer** | **Basal Lamina** | *p-value* |
| ***As*** | | | | |
| 18-20 hpi | <BL,CL | >BM | >BM | 1,70E-18 |
| 21-23 hpi | <CL | >BM | ns | 1,21E-07 |
| 24-25 hpi | <BL,CL | >BM | >BM | 2,30E-09 |
| ***Ag*** | | | | |
| 18-20 hpi | <BL,CL | >BM | >BM | 4,20E-13 |
| 21-23 hpi | <CL | >BM | ns | 2,40E-11 |
| 24-25 hpi | <BL,CL | >BM | >BM | 3,00E-04 |
| $Ag^{TEP1KD}$ | | | | |
| 18-20 hpi | <BL,CL | >BM | >BM | 2,50E-07 |
| 21-23 hpi | <CL | >BM | ns | 1,00E-04 |
| 24-25 hpi | <BL,CL | >BM | >BM | 7,94E-11 |

**Table S10.** Number of dextran-positive cells in *A. gambiae* and *A. stephensi* mosquitoes at 18-25 h post infection.

| | ***A. stephensi*** | ***A. gambiae*** |
|---|---|---|
| Records analyzed (n) | 11 | 23 |
| Dextran positive cells (n) | 17 | 52 |
| Dextran positive cells containing a parasite (n) | 12 | 20 |
| Dextran positive cells with an ookinete, % | **71** | **39** |
| -within the Blood meal, % | **71** | **50** |
| -within the Cell layer, % | **0** | **31** |

**Table S11.** Summary of phenotypes *A. stephensi* (*As*), *A. gambiae* (*Ag*) and *A. gambiae* with silenced *TEP1* (*Ag$^{TEP1KD}$*).

| | ***As*** | ***Ag*** | ***Ag$^{TEP1KD}$*** |
|---|---|---|---|
| **Infection level** | high | low | high |
| **Proportion of parasites in Blood meal, %** | ~70 | ~70 | ~20 |
| **Ookinete speed in Blood meal, um/min** | 8.2 | 3.5 | -- |
| **Proportion of parasites in Cell layer, %** | ~20 | ~20 | ~50 |
| **Ookinete speed in Cell layer, um/min** | 0.4 | 1.8 | |
| **proportion of intercellular ookinetes, %** | ~50 | ~25 | ~30 |
| **Dextran filled cells contain a parasite, %** | ~70 | ~40 | |
| **Proportion of parasites in Basal lamina, %** | ~10 | ~10 | ~20 |
| **Ookinete speed in Basal lamina, um/min** | 0.3 | 0.5 | -- |
